# Repression of Transcription Factor AP-2 Alpha by Peroxisome Proliferator Activated Receptor Gamma Reveals a Novel Transcriptional Circuit in basal-squamous Bladder Cancer

**DOI:** 10.1101/401307

**Authors:** Hironobu Yamashita, Yuka Imamura Kawasawa, Lauren Shuman, Zongyu Zheng, Truc Tran, Vonn Walter, Joshua I. Warrick, Hikmat Al-Ahmadie, Matthew Kaag, Pak Kin Wong, Jay D. Raman, David J. DeGraff

## Abstract

The discovery of bladder cancer transcriptional subtypes provides an opportunity to identify high risk patients, and tailor disease management. Recent studies suggest tumor heterogeneity contributes to “plasticity” of molecular subtype during progression and following treatment. Nonetheless, the transcriptional drivers of the aggressive basal-squamous subtype remain unidentified. As PPARγ has been repeatedly implicated in the luminal subtype of bladder cancer, we hypothesized inactivation of this transcriptional master regulator during progression results in increased expression of basal-squamous specific transcription factors (TFs) which act to drive aggressive behavior. We initiated a pharmacologic and RNA-seq-based screen to identify PPARγ-repressed, basal-squamous specific TFs. Hierarchical clustering of RNA-seq data following treatment of a panel of human bladder cancer cell lines with a PPARγ agonist identified a number of TFs regulated by PPARγ activation, several of which are implicated in urothelial and squamous differentiation. One PPARγ-repressed TF implicated in squamous differentiation identified is Transcription Factor Activating Protein 2 alpha (TFAP2A). We show TFAP2A and its paralog TFAP2C are overexpressed in basal-squamous bladder cancer and in squamous areas of cystectomy samples, and that overexpression is associated with increased lymph node metastasis and distant recurrence, respectively. Biochemical analysis confirmed the ability of PPARγ activation to repress TFAP2A, while PPARγ antagonist studies indicate the requirement of a functional receptor. *In vivo* tissue recombination studies show TFAP2A and TFAP2C promote tumor growth in line with the aggressive nature of basal-squamous bladder cancer. Our findings suggest PPARγ inactivation, as well as TFAP2A and TFAP2C overexpression cooperate with other TFs to promote the basal-squamous transition.

## Introduction

While the most commonly diagnosed type of bladder cancer (BC) is morphologically defined as urothelial carcinoma, the existence of morphologic variants of BC and their association with clinical outcomes has been recognized for decades. Molecular studies show that variant morphologies in BC exhibit unique gene expression patterns, which may contribute to differing oncologic outcomes in these patients (1,2). Moreover, recent studies have identified a striking degree of intra-tumoral, morphologic and molecular heterogeneity in advanced BC (3). If advanced BC with intratumoral heterogeneity is largely clonal in nature, the fact that the vast majority of carcinoma *in situ* (considered the precursor for the majority of advanced BC) lesions are luminal (4) strongly suggests molecular subtype is “plastic” and can evolve over time. This perspective is substantiated by the fact that areas of variant morphology exhibit significant differences in gene expression subtype within a single tumor, yet harbor a large number of identical genetic alterations (5).

While the exact temporal sequence of genetic alterations in BC and how these alterations directly contribute to tumor heterogeneity is unknown, several lines of evidence implicate the steroid hormone receptor peroxisome proliferator active receptor gamma (PPARγ) in morphologic and molecular plasticity. For example, activation of this nuclear hormone receptor has been shown to oppose squamous differentiation (SqD) *in vitro* (6), while inactivation of both PPARγ and PTEN expression drive squamous changes *in vivo* (7). Moreover, PPARγ is amplified at the genomic level in the luminal BC subtype where it is consistently overexpressed (8-10), and activation of PPARγ cooperates with overexpression of FOXA1 and GATA3 to “reprogram” the basal-squamous cell line 5637 to exhibit a luminal expression pattern (11). While these observations suggest PPARγ is a master regulator of luminal BC cell fate, as well as a potential therapeutic target (8,12-14), the transcriptional mediators of basal-squamous BC remain unidentified.

Accordingly, we hypothesized PPARγ actively represses transcription factors (TFs) that drive basal-squamous gene expression in human BC, and by extension, inactivation of PPARγ drives expansion of basal-squamous clones by upregulating these TFs. We tested the initial component of this hypothesis in the current study by performing a pharmacologic and RNA-seq based screen to identify PPARγ-repressed TFs operative in driving the basal-squamous subtype. In doing so, we provide the first evidence identifying members of the Transcription Factor Activator Protein 2 (TFAP2) family as markers of basal-squamous BC that play a direct role in mediating the phenotype of this aggressive subtype of disease.

## Materials and Methods

### Cell culture

The UMUC1 cell line was purchased from the European Collection of Authenticated Culture Collection (Salisbury, UK). Additional lines were purchased from the American Type Culture Collection (ATCC; Manassas, VA). Cell origin was confirmed by short tandem repeat (STR) analysis (11). Culture medium was used as follows: McCoy’s Modified Medium with 10% Fetal bovine serum (FBS; Sigma; St. Louis, MO) (RT4 and T24), Minimal Essential Medium (MEM) with 10% FBS (UMUC1 and UMUC3), MEM with Non-essential amino acids (NEAA), sodium pyruvate and 10% FBS (SCaBER, TCCSUP, HT-1376, HT-1197), RPMI1640 with 10% FBS (SW780 and 5637). The Lenti-X 293T cell line (Takara, Mountain View, CA) was maintained in Dulbecco’s Modified Eagle’s Medium (DMEM) with high glucose, L-glutamine with 10% Tet System Approved FBS (Takara). All cell culture medium was purchased from Thermo Fisher (Waltham, MA) and supplemented with penicillin/streptomycin.

### PPARγ agonist and antagonist treatment

One day before transfection, 5637, UMUC1, SW780 BC cells were plated (4×10^5^ cells/well) in 6 well plates (Corning Inc., Corning, NY) in complete culture medium containing 10% FBS and allowed to attach overnight. On the following day, culture medium were removed and attached cells were washed one time with serum-free medium (serum-free RPMI1640 medium for 5637 and SW780 cells, serum-free MEM/EBSS medium for UMUC1 cells) and then replaced with serum-free medium and incubated for an additional 24 hours. After 24 hours of serum starvation, medium was replaced with serum free medium containing either Dimethyl sulfoxide (DMSO; Sigma) vehicle control or rosiglitazone (TZD; 1μM; TOCRIS; Bristol UK). Cells were treated in the presence of DMSO or TZD for 48 additional hours, and RNA and protein were harvested via routine approaches as described below in the pertinent methods sections. For experiments utilizing the PPARγ antagonist GW9662 (TOCRIS), cells were pretreated with GW9662 at a final concentration of 1 or 5 μM for 1 hour before the addition of TZD.

### RNA-sequencing and computational analysis

cDNA libraries were prepared using the NEXTflex™ Illumina Rapid Directional RNA-Seq Library Prep Kit (Bio Scientific) as per the manufacturer’s instructions. Denatured libraries were diluted to 10 pM by pre-chilled hybridization buffer and loaded onto a TruSeq SR v3 flow cells on an Illumina HiSeq 2500 for 50 cycles using a single-read recipe (TrueSeq SBS Kit v3) and run for 50 cycles using a single-read recipe according to the manufacturer’s instructions. De-multiplexed and adapter-trimmed sequencing reads were generated using Illumina bcl2fastq (released version 2.18.0.12) allowing no mismatches in the index read. The sequencing reads were subjected to quality filtering used FASTX-Toolkit (http://hannonlab.cshl.edu/fastx_toolkit) to keep only reads that have at least 80% of bases with a quality score of 20 or more (conducted by fastq_quality_filter function) and reads left with > 10 bases after being trimmed with reads with a quality score of < 20 (conducted by fastq_quality_trimmer function). Filtered reads were mapped to the human reference genome (GRCh38) using TopHat (version 2.0.9) (15) supplied by Ensembl annotation file; GRCh38.78.gtf. After normalization was performed via the median of the geometric means of fragment counts across all libraries, differential expression was determined using the Cuffdiff tool which is available in Cufflinks version 2.2.1 (16). All genes passing FDR criteria of 0.05 were considered differentially expressed genes. Venn diagrams and heatmaps were generated using limma in R. GO analysis was performed using DAVID Bioinformatics Resources 6.8 (https://david.ncifcrf.gov). For computational analysis of TCGA data for TFAP2A and TFAP2C expression, RNA-seq expression data was obtained from the Genomic Data Commons (https://portal.gdc.cancer.gov/). Expression was log2 normalized and clustered with the genes listed using the pheatmap package in R. pheatmap: Pretty Heatmaps. R package version 1.0.10 (https://CRAN.R-project.org/package=pheatmap) and correlation distance. Annotation data including expression subtype were obtained from the supplemental data available with the TCGA bladder cancer project (9).

### RNA extraction and quantitative real time PCR (qRT-PCR)

Total RNA was extracted using the RNeasy approach (Qiagen; Hilden, Germany) according to manufacturer protocol. For cDNA synthesis, reverse transcription was performed using M-MLV reverse transcriptase (Thermo fisher) via manufacturer instructions. qRT-PCR was performed using QuantaStudio7 Real-Time PCR System (Applied Biosystems; Foster City, CA). Taqman probes used in this study were as follows. TFAP2A (Hs01029413_m1), TFAP2C (Hs00231476_m1), FABP4 (Hs01086177_m1). Relative gene expression change was calculated by deltadeltaCt method. 18S ribosomal RNA was used as an endogenous reference.

### Western blotting

Cell protein extracts were prepared with radioimmunoprecipitation assay buffer (Thermo Fisher) containing 25mM Tris-HCl (pH7.6), 150 mM NaCl, 1 % NP-40, 1% sodium deoxycholate, 1 % SDS including 1x proteinase inhibitor for 15 minutes on ice. Lysates were sonicated and centrifuged at 15,000 rpm in an Eppendorf (Hamburg, Germany) 5424R refrigerated centrifuge for 10 minutes and supernatants were collected. A total of 10 μg of lysate was used with 1x lithium dodecyl sulfate (LDS) sample buffer containing 10% 2-mercaptoethanol (Sigma). Gel electrophoresis was performed on 4-12% gradient Bis-Tris NuPAGE gels (Thermo Fisher) and proteins were subsequently transferred to Immobilon-P polyvinylidene fluoride (PVDF) membrane (Millipore; Burlington MA) using a PierceG2 Fast Blotter (Thermo Fisher). Membranes were blocked with 5% skim milk in Tris buffered saline containing 0.1% Tween-20 (Bio-Rad; Hercules CA) (TBST) for 1 hour at room temperature. After blocking, primary antibodies (see Supplementary Table S1) were diluted with 5% skim milk in TBST and incubated overnight at 4°C and washed three times for 10 minutes in TBST. Secondary antibodies (anti-rabbit horseradish peroxidase (HRP) conjugated IgG (1:2000) (GE Healthcare, Chicago IL), anti-mouse HRP conjugated IgG (1:2000) (GE Healthcare, Chicago IL), and anti-goat HRP conjugated IgG (1:2000) (HAF109, R&D Systems)) were diluted with 5% skim milk in TBST and incubated for 1 hour at room temperature. Following incubation with secondary antibodies, membranes were washed three times for 10 minutes with TBST, and membranes were exposed to CL-X Posure^TM^ films (Thermos Fisher) following incubation with ECL plus Western Blotting Detection Reagents (GE Healthcare).

### Plasmid construction

For construction of pLVX IRES-TFAP2A-DDK plasmid, cDNA was amplified by Accuprime Super Mix (Invitrogen; Carlsbad, CA) using UltimateORF TFAP2A/AP2alpha (Thermo Fisher, IOH46467) as a template. The following primers were used in this reaction. Forward (5’-TCTGAATTCACCATGCTTTGGAAATTGACGGATAATATC-3’),Reverse (5’GCTCGAGTTAAACCTTATCGTCGTCATCCTTGTAATCCAGCTTTCTGTGCTTCTCCTCTTTGTCACTGCTTTTG-3’). PCR products were digested with *EcoRI* and *XhoI* (New England Biolabs; Ipswich, MA) and ligated into the pLVX-IRES-Neo^R^ plasmid. pLVX-IRES-TFAP2A-DDK plasmid was analyzed by sequencing. In order to establish pLVX-EF1alpha-TFAP2C-V5pLenti6.3/V5-TFAP2C was generated first by mixing 1μl of Ultimate ORF TFAP2C (IOH28749, Thermo Fisher) and 1μl of pLenti6.3/V5-DEST (Thermo Fisher) in TE buffer (pH8.0) and incubating with 2μl of LR Clonase^TM^ II at 25 °C for 1 hour. This mixture was subsequently incubated with 1μl of ProteinaseK (Sigma) and transformed into chemically competent *E. coli* cells. Positive clones were sequenced using primers, CMV-forward (5’-CGCAAATGGGCGGTAGGCGTG-3’) and V5-reverse (5’-ACCGAGGAGAGGGTTAGGGAT-3’). Secondly, site-directed mutagenesis was performed to exchange TAG into CAGA in-frame to ablate the stop codon before the V5 tag sequence in the pLenti6.3/V5-TFAP2C plasmid using the QuickChange^®^ Site-Directed Mutagenesis kit (Thermo Fisher). Using the pLenti6.3/V5-TFAP2C plasmid as a template, a PCR product was amplified using the following primers: Forward (5’-GAGAAACACAGGAAACAGAACCCAGCTTTCTTGTACA-3’) and Reverse (5’-TGTACAAGAAAGCTGGGTTCTGTTTCCTGTGTTTCTC-3’) in the presence of 5% DMSO. Amplified products were digested with *DpnI* and transformed into XL1-blue competent cells. Transformants were sequenced using primers of CMV-forward (5’-CGCAAATGGGCGGTAGGCGTG-3’) and V5-reverse (5’-ACCGAGGAGAGGGTTAGGGAT-3’). To construct pLVX-EF1alpha-IRES-TFAP2C-V5 plasmid, the insert containing TFAP2C and V5 tag was amplified by Platinum Taq polymerase (Thermo Fisher) based on pLenti6.3/V5-TFAP2CMut plasmid generated by mutagenesis above as a template. Primers were used Forward (5’-TTTACTAGTATGTTGTGGAAAATAACCGATAATGTC-3’) and Reverse (5’-TTAGGATCCCTAACCGGTACGCGTAGAATCGAG-3’). The amplified PCR product was digested with *SpeI* and *BamHI* and was ligated into pLVX-EF1alpha-IRES-puro^r^ (Clonetech).

### Generation of stable cell lines

One day before transfection, 4×10^6^ Lenti-X 293T cells in 8ml of medium were seeded into 100 mm dishes. 7 μg of pLVX-IRES-TFAP2A or pLVX-EF1alpha-IRES-TFAP2C plasmid were diluted with 600 μl of sterile water and mixed with Lenti-X Packaging Single Shots (VSV-G) (Clontech). This mixture was added to medium in a 100 mm dish and incubated for 4 hours at 37 °C. After 4 hours, an additional 6 ml fresh medium was added and incubated for 48 hours. After 48 hours, medium was collected and virus titer was measured by Lenti-X GoStix^TM^ (Clontech). Collected medium containing Lentivirus was filtered through a 0.45 μm polyethersulfone membrane (GE Healthcare, Chicago IL). For lentivirus infection, T24 or UMUC3 cells were seeded onto 6 well plates to reach approximately 70% confluency. Medium was replaced with 2 ml of virus medium as prepared above with polybrene. After 48 hours incubation, medium was removed and replaced with medium containing 2 mg/ml G418 (for pLVX-IRES-TFAP2A) or 1 μg/ml puromycin (for pLVX-EF1alpha-IRES-TFAP2C).

### siRNA transfection

SiRNA transfection was performed using a reverse transfection method with lipofectamine 3000 (Thermo Fisher). SiRNAs for non-targeting siRNA (Scrambled; D-001810-01-05), TFAP2A siRNA (L-006348-02-0005) and TFAP2C siRNA (L-005238-00) were obtained from Dharmacon (Lafayette, CO). siRNA and lipofectamine 3000 were added to separate aliquots of OPTi-MEM medium, respectively and incubated for 5 minutes. Subsequently, siRNA and lipofectamine 3000 in OPTi-MEM were mixed and incubated for 20 minutes. After incubation, siRNA-lipofectamine 3000 complex was added to 2 ml cell suspension containing 4×10^5^ target cells per well in a 6 well plate.

### Immunohistochemistry of human bladder cancer tissue

Immunohistochemistry (IHC) on our previously described human BC TMA (17) was performed via established methods (18). Primary antibodies are referenced in Supplementary Table S1. Nuclear expression was quantified by Allred score, which combines a measure of expression intensity (range 0 to 3, no expression to highest expression) to a measure of expression area (range 0 to 5, no expression to diffuse expression) to give a score ranging form 0 to 8 (19).

### Migration and Invasion assays

Falcon 8.0 μm transparent PET membrane cell culture inserts were used for migration assays and Matrigel Transwell inserts (Corning) was used for invasion assays. Briefly, 750μl of medium containing 10% FBS was added to the space between cell culture insert or a Matrigel Transwell insert (Corning) and the well within a 24 well plate. A total of 1×10^5^ cells were suspended in 500μl of serum free medium and added to the transwell insert. T24 and UMUC3 cells were incubated for 12 hours (migration), or 24 hours (invasion). SCaBER cells were incubated for 12 hours (migration), or 15 hours (invasion). After incubation, migrated and invaded cells were stained with 0.5 % crystal violet (Sigma) in 20% methanol (Sigma) for 20 minutes, and residual cells that had not moved through the transwell were removed by gentle swabbing with a Q-tip (Unilever, Trumbull, CT). Cell numbers were counted in microscopic field (x100 magnification). All experiments were repeated at least 3 times in triplicate.

### Tissue recombination xenografting

All animal experiments were performed in accordance with approved protocols from the Institutional Animal Care and Use Committee (IACUC) of Pennsylvania State University. Isolation of embryonic bladder mesenchyme (eBLM), preparation of tissue recombinants, and kidney capsule surgeries were performed as described previously (7,11,20). Pregnant rats (Harlan; Indianapolis, IN) were sacrificed at embryonic day 16 (plug day = 0). Embryos were isolated as previously described, and bladders were microdissected under a dissecting microscope (Olympus SZX7; Center Valley, PA) from isolated embryos, and embryonic bladders were separated from the urogenital sinus at the bladder neck and the attached ureters carefully dissected using Vanna spring scissors (Fine Science Tools; Foster City, CA). Followng incubation in calcium and magnesium-free Hanks’ saline (Thermo Fisher) containing 25mM EDTA (Sigma), embryonic bladder mesenchyme (eBLM) and urothelium were separated manually under microscopic examination using two 25 gauge needles connected to a 1cc syringe, leaving the eBLM behind as a “shell.” UMUC3 and T24 (1× 10^5^) cells overexpressing TFAP2A or TFAP2C were re-suspended in 50 μl of a 3:1 ratio of rat tail collagen and setting solution, and were plated in 10cm dishes. Following the insertion of 1 eBLM per aliquot, tissue recombinants were placed at 37°C to promote solidification. McCoy’s modified medium (Thermo Fisher) containing 10 % FBS was then applied to solidified grafts and incubated overnight. The following day, tissue recombinants were placed under the kidney capsule of the left kidney of 5 SCID mice (Jackson Laboratories; Bar Harbor, MA), resulting in a total of 10 grafts for each cell line. Four weeks following implantation, mice bearing UMUC3 cells were sacrificed, whereas mice bearing T24 cells were sacrificed 8 weeks after surgery, respectively. Tumor radius was measured through cellSens software on an Olympus CX41 microscope, and tumor volumes were approximated by the calculation tumor volume = π (r^2^).

### Statistical analysis

Statistical analysis was performed using SAS (SAS Institute Incoprotated; Cary, NC) and GraphPad Prism6 (GraphPad Software; San Diego, CA). *In vitro* data were analyzed using parametric tests (Student’s t-test and analysis of variance (ANOVA) with Tukey’s *post hoc* correction for multiple comparison), while *in vivo* and clinical data was analyzed using non-parametric (Wilcoxon rank sum test and Kruskal-Wallis H test) approaches. Chi-square tests were used to test for associations between TFAP2A and TFAP2C expression with lymph node metastasis and distant recurrence. *In vitro results* are expressed as the mean ± S.D. p<0.05 was considered as a statistically significant.

## Results

### PPARγ is a master regulator of luminal gene expression in bladder cancer cells

In a previous study (11), we reported that overexpression of FOXA1 and GATA3 cooperated with PPARγ activation to “reprogram” a basal-squamous BC cell line (5637), resulting in the activation of a luminal molecular signature (see diagram, Figure 1A). Although no single factor in these experiments was able to reprogram 5637, the largest shift in gene expression in these experiments followed PPARγ activation with the agonist rosiglitazone (TZD). Based on this observation, we hypothesized the existence of a set of basal-squamous-specific TFs that are repressed in the presence of active PPARγ signaling. In an effort to identify PPARγ-regulated TFs, we first screened 6 common, PPARγ positive (11) BC cell lines (UMUC1, SW780, SCaBER, 5637, HT1376 and HT1197) for responsiveness to the PPARγ agonist TZD. Western blotting for the putative PPARγ response gene Fatty Acid Binding Protein 4 (FABP4) following 48 hours of TZD treatment confirmed 5637 responsiveness to PPARγ activation, and additionally identified the BC cell lines UMUC1 and SW780 as PPARγ-responsive (Figure 1B). These 3 cell lines were subsequently used for RNA-Seq studies to identify PPARγ-regulated TFs (see Figure 1C for experimental design). While this approach identified a number of cell line-specific TZD-regulated genes following treatment, we additionally identified a total of 26 and 10 genes coordinately upregulated and downregulated amongst 5637, UMUC1 and SW780 following PPARγ activation, respectively (Figure 1D; see Supplementary Table S2). Consistent with previous reports indicating a central role for PPARγ in the BC cell autonomous regulation of cytokine production (8), Gene Ontology (GO) enrichment analysis (Figure 1E) identified several immune related pathways as being associated with genes upregulated following TZD treatment. While GO enrhichment analysis of genes which decreased following TZD treatment identified alterations in several immune pathways in SW780, pathways associated with TZD treatment of UMUC1 and 5637 were associated largely with cell growth and gene expression (see Supplementary figure S1). As maintenance of a given molecular subtype in BC most likely results from the activity of a small subset of transcriptional master regulators (21), we further examined our RNA-seq results in an effort to identify altered expression of key TFs implicated in urothelial differentiation following PPARγ activation. RNA-seq data were analyzed using a gene set consisting of 20 TFs implicated in either urothelial differentiation or basal-squamous BC. Hierarchical clustering of RNA-seq data from all 3 cell lines predictably identified 3 main groups (one for each cell line), with subgroups apparently being driven by DMSO or TZD treatment (Figure 1F). However, individual hierarchical clustering of RNA-seq data from UMUC1 (Figure 1G), 5637 (Figure 1H) and SW780 (Figure 1I) identified Transcription Factor AP-2 alpha (TFAP2A) and Brother Of the Regulator of Imprinted Sites (BORIS/CTCF) as TZD-repressed transcription factors in all 3 cell lines, while the pluripotency factor Kruppel Like Factor 4 (KLF4) expression was increased by TZD treatment in all three cell lines (Figure 1 G-I and Supplementary Figure S2). Our findings are in agreement with previous reports indicating a role for BORIS/CTCF (22) and KLF4 (23) in urothelial differentiation and BC, respectively and additionally suggest a role for TFAP2A in the emergence of a basal-squamous phenotype following PPARγ inactivation in BC.

**Figure 1.**
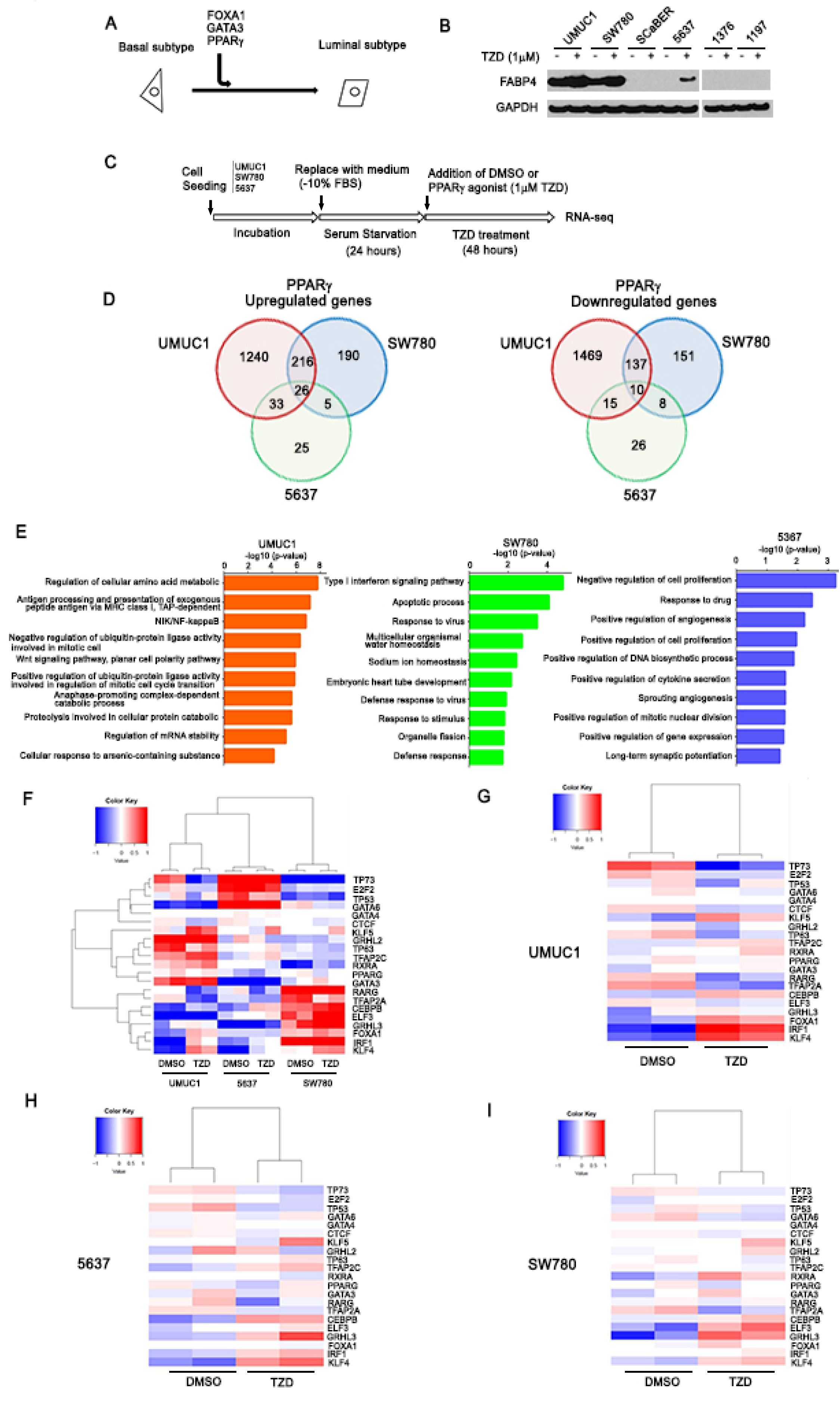
Identification of PPARγ-repressed transcription factors. **A:** Our previous study (11) shows FOXA1, GATA3 and PPARγ activation cooperate to “reprogram” a human basal BC cell line (5637) to exhibit a luminal gene expression subtype, thus providing a platform for the identification of PPARγ-regulated genes. **B:** Western blot analysis of FABP4 in UMUC1, SW780, SCaBER, 5637, HT1376, and HT1197 after 48 hours PPARγ agonist (Rosiglitazone (TZD)) treatment. **C:** Experimental design for PPARγ activation via TZD treatment in BC cells (see materials and methods). **D:** Venn diagram showing shared and unique sets of significantly upregulated (left) or downregulated (right) genes following 48 hours TZD treatment of UMUC1, SW780 and 5637 cells. **E:** GO analysis based on significantly upregulated genes following PPARγ activation. Biological process by GO analysis based on significantly upregulated genes following PPARγ activation in UMUC1, SW780, and 5637 cells. Top 10 biological process cetegories of significantly upregulated by PPARγ activation are shown. The vertical axis shows biological process cetegories, and horizontal axis shows the –log10(P-value). **F:** Heat map shows hierarchical clustering of RNA-seq data following treatment of UMUC1, SW780 and 5637 with vehicle control (DMSO) or TZD for 48 hours. **G:** Heat map shows hierarchical clustering for UMUC1 alone after drug treatment. **H:** Heat map shows hierarchical clustering for 5637 alone after drug treatment. **I:** Heat map shows hierarchical clustering for SW780 alone after drug treatment. Genes for clustering analysis based on previous studies suggesting a role for urothelial differentiation and/or differential expression in BC molecular subtypes. Expression values are median centered by gene in all heatmap displays.

### PPARγ signaling represses TFAP2A expression in bladder cancer cells

We next investigated the regulatory relationship between PPARγ and TFAP2A. Treatment of serum-starved 5637, UMUC1 and SW780 with 1 µM TZD for 48 hours followed by Q-RT-PCR for FABP4 confirmed responsiveness of these lines to PPARγ activation (Figure 2A). In addition, Q-RT-PCR and western blotting confirmed the ability of PPARγ activation to repress TFAP2A expression at the mRNA (Figure 2B) and protein (Figure 2C) levels, respectively. To confirm a role for a functional PPARγ receptor, we performed identical TZD treatments of 5637, UMUC1 and SW780 alone and in conjunction with the PPARγ antagonist GW9666. Q-RT-PCR results show that while TZD treatment significantly increased FABP4 expression and decreased TFAP2A expression, these significant changes were abolished in the presence of GW9662 co-treatment (Figure 3A and 3B). In addition, while TZD treatment significantly reduced TFAP2A protein levels, this was prevented following co-treatment with GW9662 treatment (Figure 3C). These results suggests that TZD-induced repression of TFAP2A mRNA and protein requires a functional PPARγ receptor in BC cells.

**Figure 2.**
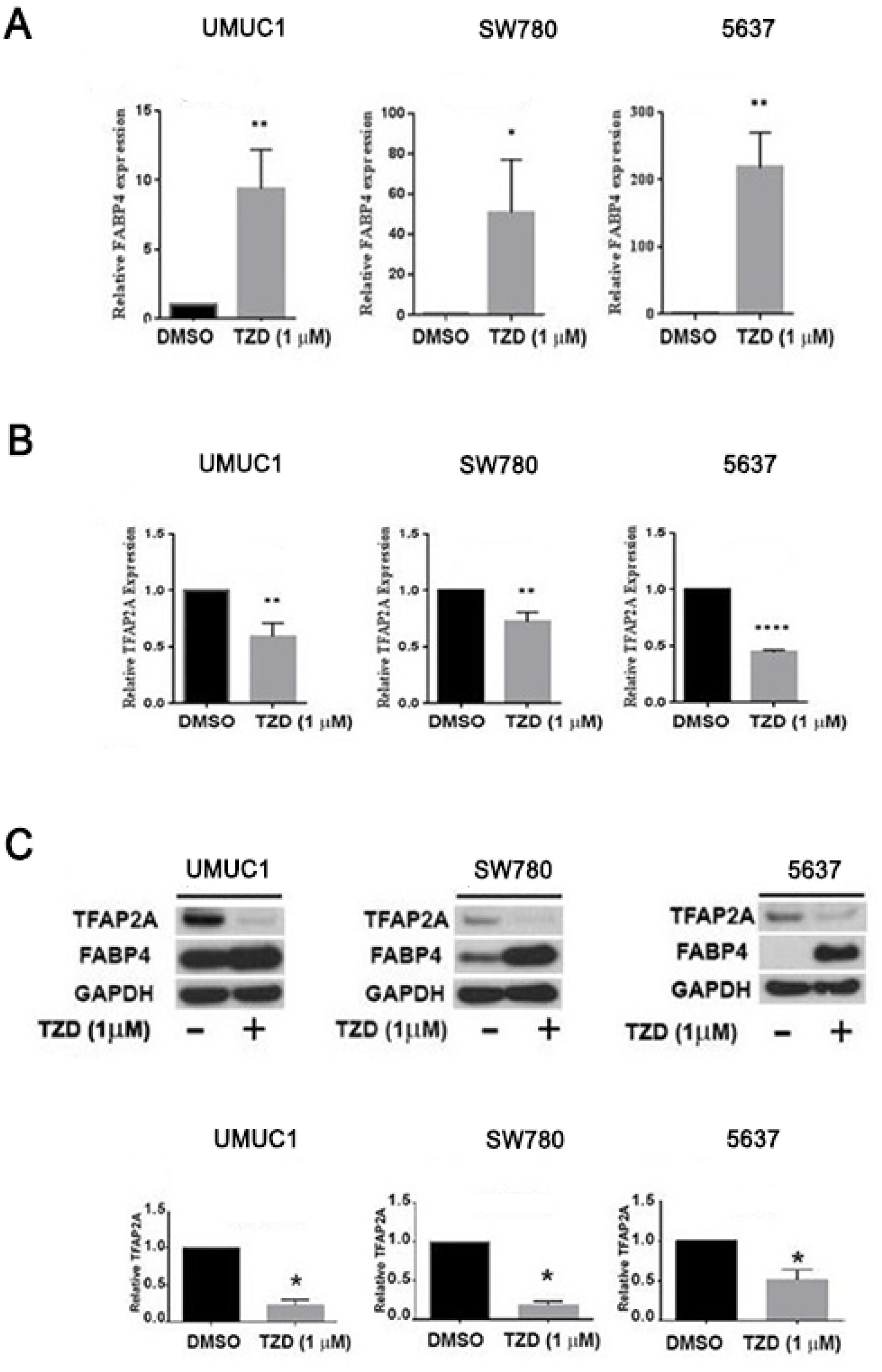
PPARγ activation represses TFAP2A expression in human bladder cancer cells. **A:** q-RT-PCR analysis of FABP4 expression levels in UMUC1, SW780 and 5637 cells after 48 hours PPARγ agonist treatment. FABP4 expression was normalized by GAPDH, internal control. * p<0.05, ** p<0.01, (Student’s *t* test). **B:** q-RT-PCR analysis of TFAP2A expression levels in UMUC1, SW780 and 5637 cells after 48 hours PPARγ agonist treatment. TFAP2A expression was normalized by GAPDH, internal control. ** p<0.01, **** p<0.0001 (Student’s *t* test). **C:** Western blot analysis of TFAP2A protein expression level in UMUC1, SW780, 5637 cells after 48 hours PPARγ agonist treatment. Densitometric analysis of western blot of TFAP2A expression (below). * p<0.05 (Student’s *t* test). In A, B, C relative expression levels of FABP4 and TFAP2A mRNA and or/protein after PPARγ agonist treatment are represented compared to that of DMSO treatment.

**Figure 3.**
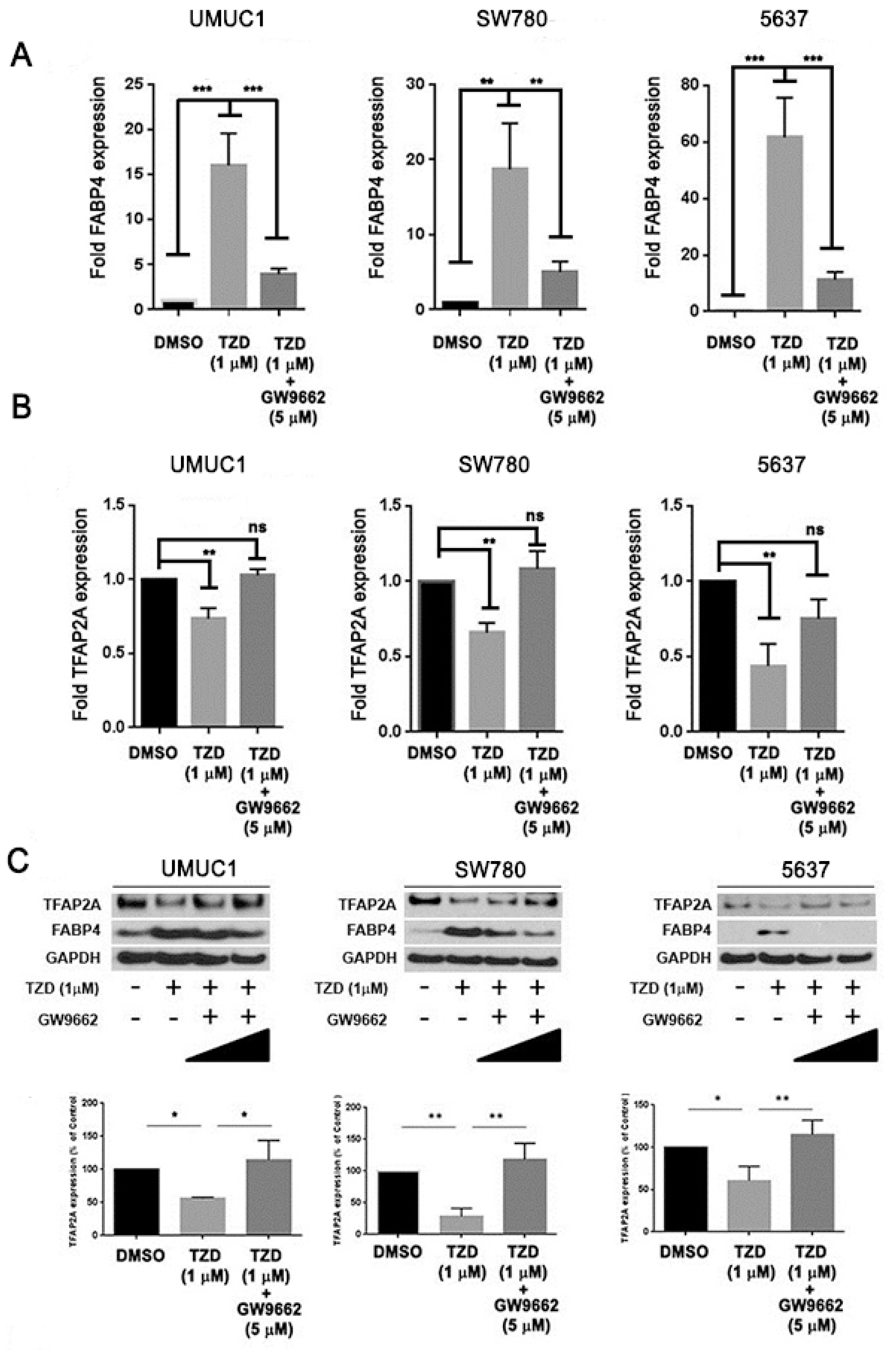
Repression of TFAP2A expression via PPARγ is dependent on a functional receptor. **A and B:** q-RT-PCR analysis of FABP4 (A) and TFAP2A (B) expression levels in UMUC1, SW780 and 5637 cells after PPARγ agonist (1μM) treatment alone or in the presence of the PPARγ antagonist (5 μM), GW9662. The putative PPARγ-regulated gene, FABP4 was used as positive control for drug treatments. Relative expression levels of TFAP2A and FABP4 after PPARγ treatment is represented relative to that of DMSO treatment. ** p<0.01, *** p<0.001, ns: not significant, one-way ANOVA with post hoc multiple comparison (Tukey). **C:** Western blotting analysis of TFAP2A protein expression levels in UMUC1, SW780 and 5637 cells after PPARγ agonist (1μM) treatment alone and in the presence (1μM and 5μM) of the PPARγ antagonist, GW9662. Densitometric analysis of western blotting of TFAP2A expression (below). TFAP2A expression was normalized by GAPDH, internal control. * p<0.05, ** p<0.01, one-way ANOVA with post hoc multiple comparison (Tukey).

### TFAP2A and TFAP2C expression are markers of basal-squamous bladder cancer

basal-squamous BC is significantly enriched for SqD, and several studies have implicated TFAP2A and its paralog TFAP2C in development and differentiation of normal squamous epithelium (24-27). Therefore, our identification of TFAP2A as a PPARγ-repressed transcriptional regulator suggested a role for members of the TFAP2 family in basal-squamous BC. We therefore utilized qRT-PCR and the delta-delta CT method to examine the expression of TFAP2A, TFAP2C and PPARγ in a panel of BC cell lines representative of luminal and basal-squamous BC, as well as a group of cell lines which does not fit under either gene expression subtype (“non-type”). While PPARγ expression did not appear to be correlated with cell line gene expression subtype (Figure 4A), TFAP2A (Figure 4B) and TFAP2C (Figure 4C) expression was upregulated in models of basal-squamous disease when compared to luminal and “non-type” cell lines. This was additionally confirmed by western blotting showing that while expression of both PPARγ isoforms (γ1 and γ2) was variable across BC cell lines, the basal-squamous cell lines SCaBER, 5637, HT1376 and HT1197 expressed relatively high TFAP2A and TFAP2C expression (Figure 4D) in comparison to luminal and “non-type” lines.

**Figure 4.**
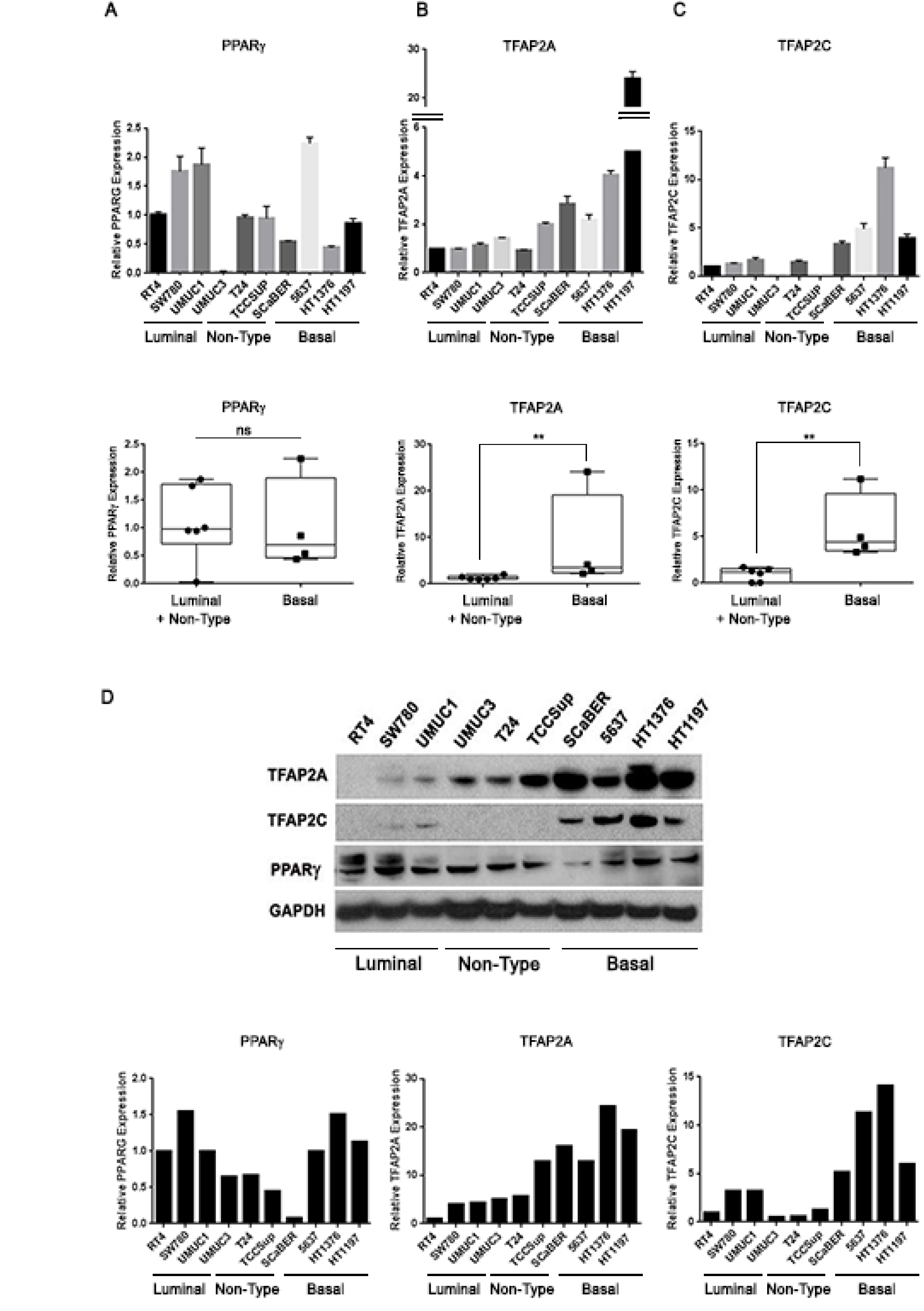
TFAP2A and TFAP2C are highly expressed in basal-squamous bladder cancer cell lines. **A-C:** q-RT-PCR analysis of mRNA expression of PPARγ (A), TFAP2A (B) and TFAP2C (C) in 10 human BC cell lines. ** p<0.01, ns: not significant, Wilcoxon rank sum test. **D:** Western blot analysis of TFAP2A, TFAP2C and PPARγ protein expression in 10 human BC cell lines. (luminal: RT4, SW780, UMUC1/ Non-type: UMUC3, T24, TCCSup/ Basal: SCaBER, 5637, HT1376, HT1197). Densitometric analysis of western blotting data is below. TFAP2A, TFAP2C and PPARγ expression was normalized by GAPDH, internal control.

While cell lines are useful *in vitro* models, they cannot faithfully represent all aspects of tumor biology. Computational analysis of publically available data through The Cancer Genome Atlas (TCGA) BC project showed TFAP2A (Figure 5A) expression is significantly enriched in basal-squamous tumors relative to other subtypes. While TFAP2C mRNA levels were not significantly correlated with molecular subtype (Figure 5B), both TFAP2A and TFAP2C clustered with basal-squamous markers including KRT5, KRT14 and TP63 (Figure 5C). In addition, immunohistochemistry performed on our previously described, in-house cohort of BC patients (17) showed invasive carcinoma with SqD expressed higher levels of TFAP2A and TFAP2C compared to invasive conventional UCC (Figure 5D-N and Supplementary Table S3; p<0.001, p=0.015, respectively, Wilcoxon rank sum test). In invasive UCC, TFAP2A expression was associated with the presence of lymph node metastasis (see Supplementary Table S4: TFAP2A: p=0.049 TFAP2C: p=0.635; Chi-square test), while increased TFAP2C expression was associated with distant recurrence (TFAP2A: p=0.137, TFAP2C: p=0.037 Chi-square test; Supplementary Table S4). These observations identify TFAP2A and TFAP2C as markers of the aggressive basal-squamous molecular subtype of human BC, as well as potentially important mediators of biologically aggressive disease.

**Figure 5.**
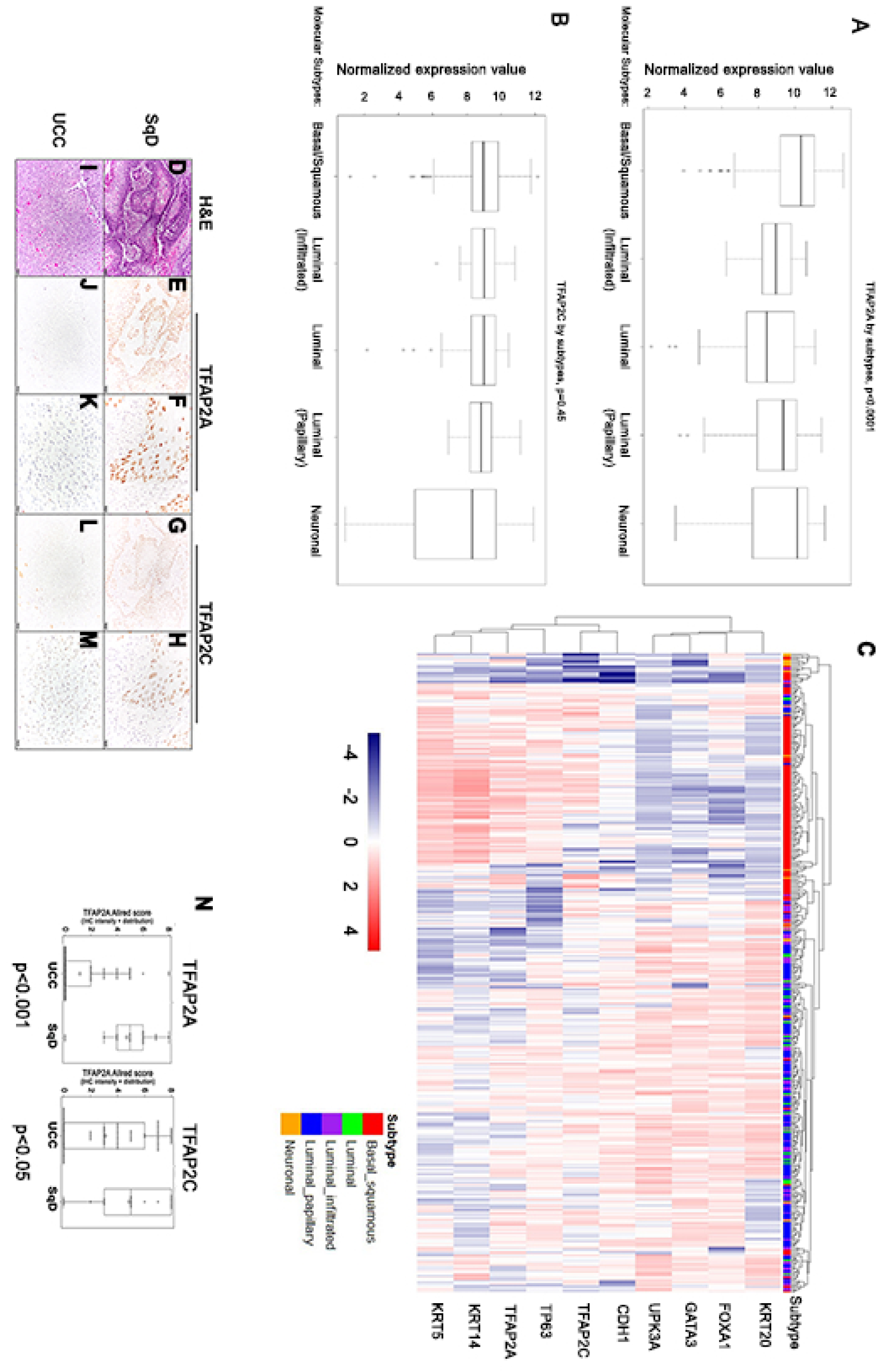
TFAP2A and TFAP2C expression is associated with basal-squamous human bladder cancer. **A and B:** Relationship between molecular subtype and expression of TFAP2A and TFAP2C was examined using data compiled through the TCGA bladder cancer study (9). While TFAP2A (A; Kruskal-Wallis H test; p <0.0001) expression was significantly elevated in the basal-squamous molecular subtype, TFAP2C (B) expression was not significantly associated with any subtype at the mRNA level. **C:** Hierarchical clustering analysis using data compiled through the same TCGA bladder cancer study shows tumors that express TFAP2A and TFAP2C cluster with tumors expressing additional markers of basal bladder cancer. **D-M:** Representative images of H&E (D, I) and IHC staining for TFAP2A (E, F, J, and K), TFAP2C (G, H, L, and M) from human BC specimens (D-H: SqD, UCC: I-M). **N**: TFAP2A (p<0.001; Wilcoxon rank sum) and TFAP2C (P<0.05; Wilcoxon rank sum) protein expression are significantly higher in SqD compared to UCC.

### TFAP2A and TFAP2C expression control the aggressive phenotype associated with basal-squamous bladder cancer cells

Because of the close association between TFAP2A and TFAP2C expression and the aggressive basal-squamous subtype, we hypothesized these transcriptional regulators may promote disease aggressiveness. Because of high TFAP2A and TFAP2C expression in SCaBER cells (See Figure 4), we performed transient knockdown of TFAP2A or TFAP2C individually or in combination in this *in vitro* model of basal-squamous disease. Western blotting (Figure 6A and B) showed we successfully decreased expression of TFAP2A and TFAP2C 24 and 48 hours post transfection. *In vitro* migration and invasion assays show that individual knockdown of TFAP2A or TFAP2C, as well as combined knockdown of TFAP2A and TFAP2C results in significantly reduced migration and invasion of SCaBER cells compared to knockdown of scramble siRNA control (Figure 6C and D). Because UMUC3 cells express relatively low levels of TFAP2A, and no detectable TFAP2C (See Figure 4D), we additionally established UMUC3 cells stably overexpressing TFAP2A (UMUC3-TFAP2A) and TFAP2C (UMUC3-TFAP2C). Western blotting (Figure 6E) and Q-RT-PCR (Figure 6F) confirmed stable overexpression of TFAP2A and TFAP2C in UMUC3 at the mRNA and protein levels, respectively. Migration and invasion assays using these stable cells showed that increased TFAP2A or TFAP2C in UMUC3 cells significantly enhanced migration and invasion (Figure 6G and H). Thus, these results suggest TFAP2A and TFAP2C regulate gene expression important for the control phenotypic aggressiveness associated with basal-squamous BC.

**Figure 6.**
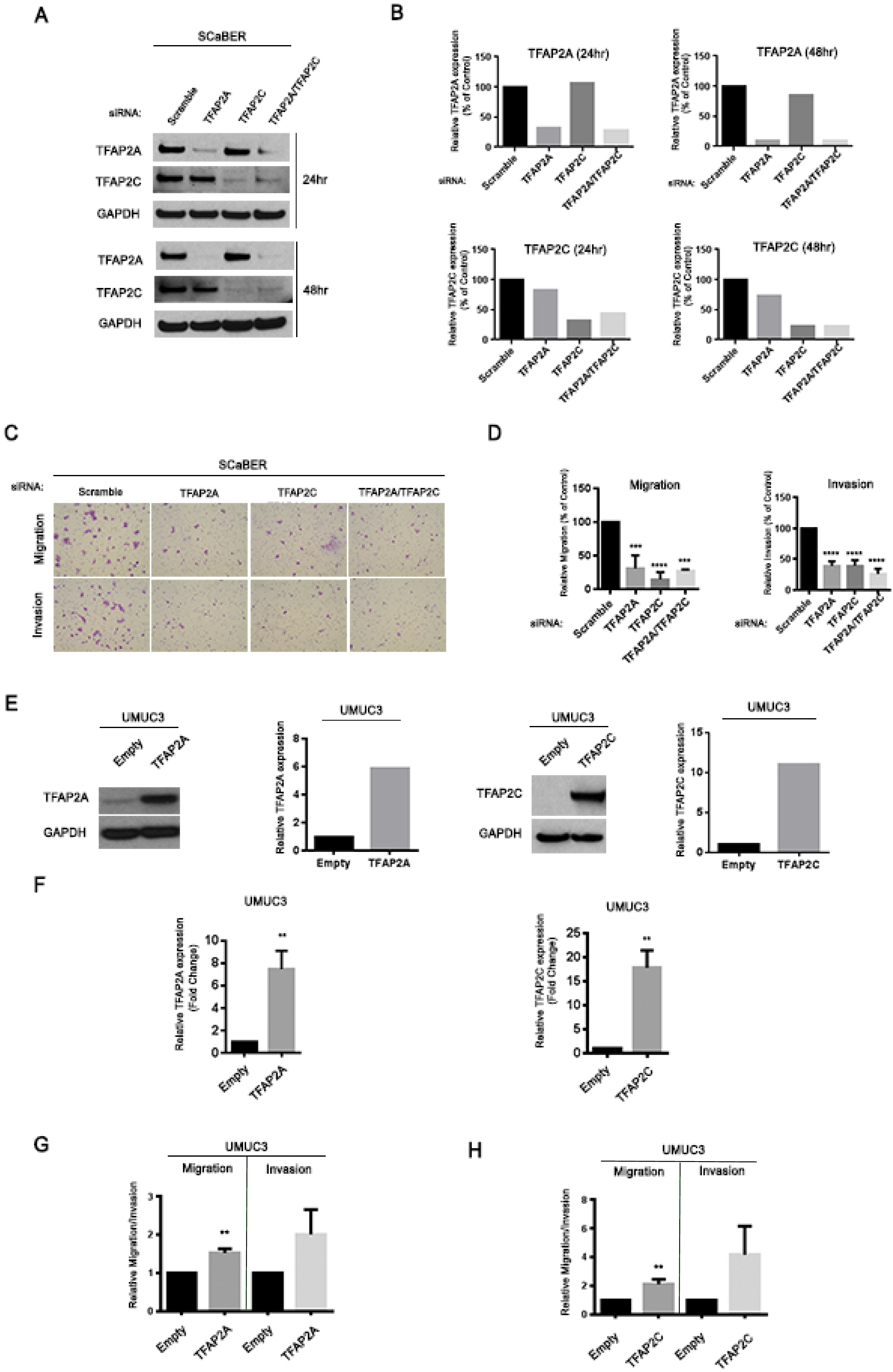
Expression of TFAP2A and TFAP2C influences BC cell migration and invasion. **A:** Western blotting analysis of TFAP2A and TFAP2C expression in SCaBER cells after siRNA transfection. SiRNAs included Scrambled (negative control), as well as constructs targeting TFAP2A and TFAP2C individually and in combination. **B:** Densitometric analysis of western blotting results for TFAP2A and TFAP2C expression for data depicted in (A). TFAP2A and TFAP2C expression was normalized to GAPDH. **C:** Representative images following migration and Invasion assays using SCaBER cells transfected with siRNA for Scrambled construct, TFAP2A, TFAP2C, and TFAP2A/TFAP2C. **D:** Relative migration and invasion of SCaBER cells transfected with siRNA. *** p<0.001, **** p<0.0001, one-way ANOVA with post hoc multiple comparison (Tukey). **E:** Western blotting analysis of TFAP2A and TFAP2C protein expression levels in UMUC3 cells overexpressing TFAP2A (left) or TFAP2C (right). Also included is densitometric analysis of western blotting data for TFAP2A and TFAP2C. **F:** q-RT-PCR analysis of TFAP2A and TFAP2C expression in UMUC3 cells overexpressing TFAP2A and TFAP2C. **G:** Migration and Invasion assay of UMUC3 stable cells overexpressing TFAP2A. **H:** Migration and Invasion assay of UMUC3 cell overexpressing TFAP2C. ** p<0.01(Student’s *t* test).

### Overexpression of TFAP2A or TFAP2C in bladder cancer cells promotes tumorigenenicity following tissue recombination xenografting

We next utilized the tissue recombination system to investigate the impact of TFAP2A and TFAP2C overexpression on tumorigenicity and the ability to drive morphologic changes, such as SqD. For these experiments, we utilized UMUC3 cells stably overexpressing TFAP2A or TFAP2C (see Figure 6E), as well as T24 cells stably overexpressing TFAP2A or TFAP2C (See Supplementary Figure S3). These cells were chosen because of their relatively low expression of TFAP2A and TFAP2C (See Figure 4). These cell lines and empty vector control cells where recombined with embryonic bladder mesenchyme, and inserted under the renal capsule of immunocompromised mice as previously described (11). Two months following implantation, T24-EV cells were weakly tumorigenic and tumor volume was not significantly increased following TFAP2A overexpression (Figure 7A and 7B; p=0.2343; quantified in 7I). However, T24 tumor volume was significantly increased following expression of TFAP2C (Figure 7C and 7D; p<0.01; quantified in 7J) 2 months following implantation. In addition, UMUC3-associated tumor volume was significantly increased following overexpression of TFAP2A (Figure 7E and 7F; p<0.05; quantified in 7K) and TFAP2C (Figure 7G and 7H; p<0.01; quantified in 7L). However, we failed to detect SqD in any of our tissue recombinants. These observations identify TFAP2A and TFAP2C overexpression as drivers of tumorigenicity, but also suggest their individual overexpression is not sufficient to drive SqD.

**Figure 7.**
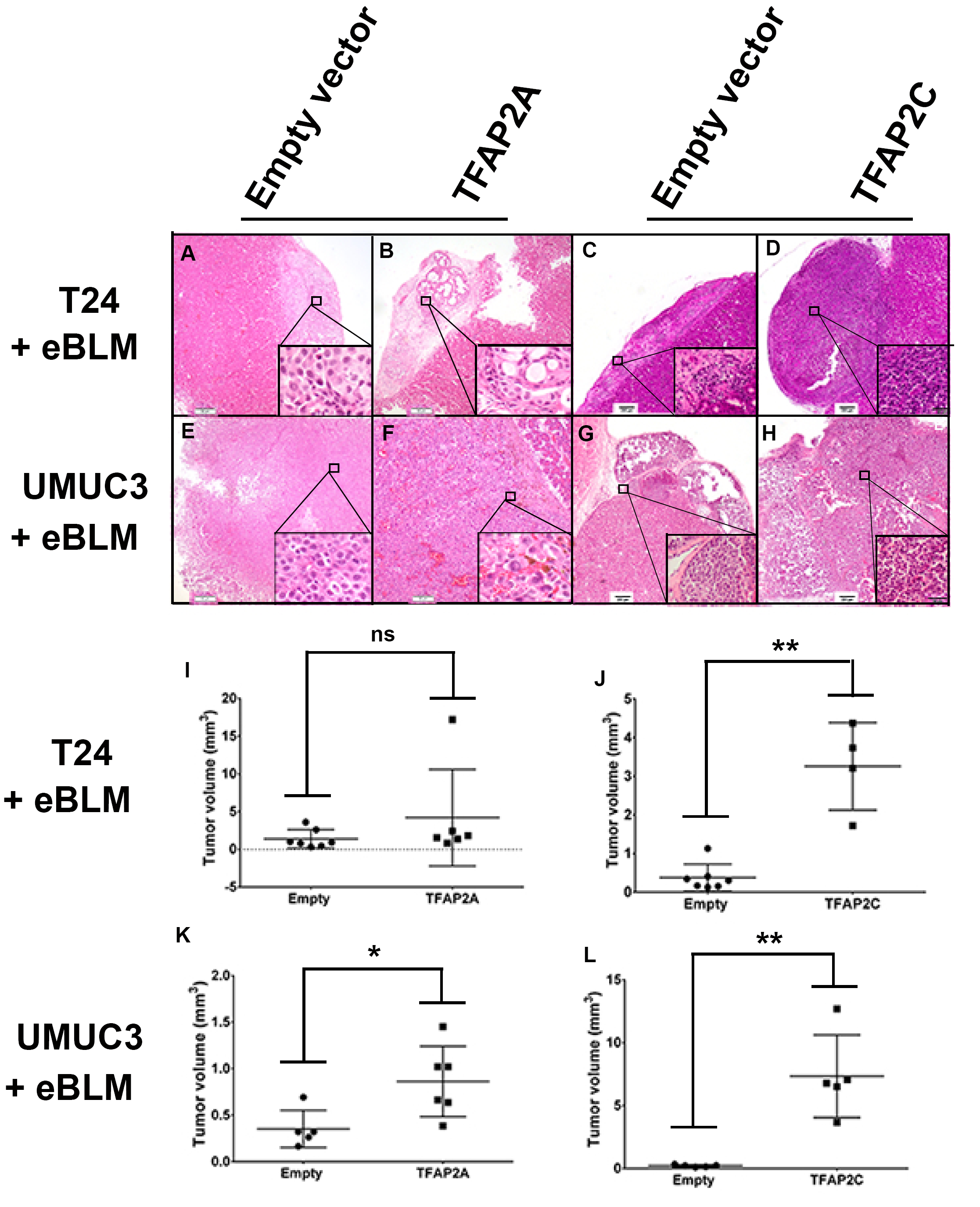
Overexpression of TFAP2A or TFAP2C in bladder cancer cell promotes tumorigenicity in tissue recombination xenografting assays. Following genetic manipulation and recombination with embryonic rat bladder mesenchyme, tissue recombinants were inserted underneath the renal capsule as described in materials and methods. **(A-D)** Hematoxylin and eosin staining of T24 cells engineered to stably express empty vector **(A)** or TFAP2A **(B),** empty vector **(C)** or TFAP2C **(D)**. **(E-H)** Hematoxylin and eosin staining of UMUC3 cells engineered to stably express empty vector **(A)** or TFAP2A **(B)**, empty vector **(C)** or TFAP2C **(D).** Overexpression of TFAP2A in T24 had no significant effect on tumor volume in the tissue recombination assay **(I)**, while overexpression of TFAP2C significantly increased tumor volume of T24 recombinants **(J).** Overexpression of TFAP2A **(K)** and TFAP2C **(L)** significantly increased tumor volume of UMUC3 recombinants. *p<0.05, ** p<0.01, ns: not significant, Wilcoxon rank-sum test.

## Discussion

Although the presence of morphologic heterogeneity in BC and its relation to clinical outcomes has been recognized for decades, a series of recent studies definitively demonstrate the existence of intratumoral molecular heterogeneity in this common malignancy (3,5), as well as the potentially plastic nature of this heterogeneity. In keeping with specific tenants of the master regulator hypothesis (21), our *in vivo* and *in vitro* experimental studies (7,11,18,20,28), as well as additional foundational studies from other groups (6,9,10,22,29-37) have identified a series of TFs apparently responsible for maintaining urothelial cell fate and establishing a luminal gene expression pattern in malignant disease. Taken together, these studies suggest PPARγ is a transcriptional master regulator of urothelial cell fate and the luminal gene expression pattern in BC. However, the positive drivers of the basal-squamous molecular subtype, as well as other molecular subtypes have remained largely unidentified.

Based on previous work suggesting PPARγ activity is critical for maintaining a luminal gene expression program (10,11), we undertook a pharmacologic screen to identify PPARγ-repressed transcription factors that act as potential master regulators of SqD and/or the basal-squamous molecular subtype (Figure 1). This screen identified TFAP2A as one such PPARγ-repressed TF that is overexpressed in human basal-squamous disease. TFAP2A belongs to the transcription factor activator protein 2 (TFAP2) family, which consists of five members (TFAP2A, 2B, 2C, 2D, 2E). Structurally, these TFs contain a helix-span-helix domain important for homo/heterodimerization, which additionally cooperates with a basic domain for DNA binding. Furthermore, a proline and glutamine rich domain serves a trans activating function. In addition to TFAP2 family members being essential for neural crest development (38) and estrogen receptor binding and subsequent long-range chromatin interactions in breast cells (39), both TFAP2A and TFAP2C are implicated in keratinocyte differentiation (27,40,41) and squamous cancers (42) independent of anatomic site. At the molecular level in normal keratinocytes, TP63 (itself implicated in the basal subtype of BC (43)) activates TFAP2C expression to promote normal skin differentiation (25,44), and cooperates with TFAP2A and TFAP2C to regulate TP63 target gene expression (26). Also, *Tfap2a* and *Tfap2c* knockout in mice produces pathologic skin disease, potentially by impacting epidermal growth factor receptor signaling which is also implicated in basal BC (41). Therefore, TFAP2A and TFAP2C play a central role in squamous-specific gene expression.

Pharmacologic approaches in our current study indicate TFAP2A regulation following TZD treatment requires functional PPARγ (Figures 2 and 3). Ligand-dependent increases in the expression of PPARγ target genes involves release of corepressor complexes including Nuclear Receptor Corepressor 1 (NCoR1) and 2 (NCoR2/SMRT) and other factors including histone deacetylases (reviewed in (45)). This process results in the recruitment of general transcription machinery and increased gene expression. However, the mechanisms of ligand-dependent repression of PPARγ target genes are less clear. The fact that TFAP2A is repressed by PPARγ activation in UMUC1, SW780 and 5637, and our observation that co-treatment with a PPARγ antagonists abrogates TZD-induced TFAP2A regulation in all 3 models suggests the existence of a general and shared mechanism. However, the exact mechanism remains to be identified. While PPARγ and other PPARs can directly interfere with the ability of other transcription factors to regulate gene expression via a process referred to as transrepression (45), it is not clear if this mechanism is responsible for ligand-dependent repression of TFAP2A by PPARγ. As PPARγ plays an important role in the BC cell autonomous regulation of cytokines and response to immunotherapy (8,46), further studies are required.

In addition to being highly expressed in *in vitro* models of basal-squamous BC (Figure 4), we additionally report here that both TFAP2A and TFAP2C are expressed at high levels in basal-squamous BC, as well as in areas of SqD (Figure 5). Importantly, the fact that TFAP2A and TFAP2C promote *in vitro* surrogate measures of aggressive behavior typically associated with basal-squamous BC (Figure 6), as well as *in vivo* tumorigenesis is in agreement with their association with lymph node metastasis and distant recurrence, and further suggests an important role for these factors in BC. In addition to the fact that SqD and the basal-squamous subtype has been associated with enhanced overall survival following neoadjuvant chemotherapy and cystectomy (47), TFAP2A was previously identified as an independent predictor of good response to cisplatin treatment in BC patients (48). While these observations further link TFAP2 family member expression to variant morphology in BC, we did not detect SqD in our tissue recombinants. However, this is not surprising as TFAP2 family members undoubtedly require the action of additional combinatorial factors (49,50) to mediate a squamous cell fate *in vivo*. Nonetheless, this study directly implicates TFAP2A and TFAP2C in the Basal molecular subtype of BC, as well as associated SqD. By additionally identifying TFAP2A as a PPARγ-repressed gene, this study has also revealed a novel transcriptional circuit which may be involved in the plasticity of molecular subtype and morphologic differentiation in this common malignancy.

## Acknowledgements

Supported in part by RSG 17-233-01-TBE from the American Cancer Society (DJD) and a Young Investigator Award from the Bladder Cancer Advocacy Network (DJD). The authors wish to acknowledge and thank Dr. Simon Hayward and Dr. Omar Franco-coronel (NorthShore Research Institute) for providing collagen and setting solution used in tissue recombination experiments. In addition, we like to acknowledge the help of Dr. Trevor Williams (University of Colorado Anschutz Medical Campus) for critical reading of the manuscript, the technical assistance of Dr. Wilson Chan and Vasy Osei Amponsa, as well as the clerical assistance of Kimberly Walker and Charity Pavlesich.

